# Spatio-temporal coherence of circadian clocks and gating of differentiation in *Anabaena* filaments

**DOI:** 10.1101/2023.02.15.528611

**Authors:** Rinat Arbel-Goren, Bareket Dassa, Anna Zhitnitsky, Ana Valladares, Antonia Herrero, Enrique Flores, Joel Stavans

## Abstract

Circadian clock arrays in multicellular filaments of the heterocyst-forming cyanobacterium *Anabaena* sp. strain PCC 7120 display remarkable spatio-temporal coherence under nitrogen-replete conditions. To shed light on the interplay between circadian clocks and the formation of developmental patterns, we followed the expression of a clock-controlled gene under nitrogen deprivation, at the level of individual cells. Our experiments showed that differentiation into heterocysts took place preferentially within a limited interval of the circadian clock cycle; that gene expression in different vegetative intervals along a developed filament was discoordinated; and that the circadian clock was active in individual heterocysts. Furthermore, *Anabaena* mutants lacking the *kaiABC* genes encoding the circadian clock core components produced heterocysts but failed in diazotrophy. Therefore, genes related to some aspect of nitrogen fixation, rather than early or mid-heterocyst differentiation genes, are likely affected by the absence of the clock. A bioinformatics analysis supports the notion that RpaA may play a role as master regulator of clock outputs in *Anabaena*, the gating of differentiation by the circadian clock and the involvement of the clock in proper diazotrophic growth. Together, these results suggest that under nitrogen deficient conditions, the functional unit in *Anabaena* is reduced from a full filament under nitrogen-rich conditions, to the vegetative cell interval between heterocysts.

## Introduction

Circadian clocks arose during evolution to enable organisms, from cyanobacteria to plants and mammals, to tune their metabolism and bioprocesses to daily light/darkness cycles on Earth, and thereby optimize their fitness (Johnson and Rust, 2021; Shultzaberger et al., 2015). Much of what is known about the mechanisms behind circadian clocks in the case of cyanobacteria has been learned from investigations of unicellular organisms, primarily *Synechococcus elongatus* strain PCC 7942 (henceforth *S. elongatus*). These investigations have firmly established that the core clock is comprised of three proteins, KaiA, KaiB and KaiC, the first two of which modulate the four phosphorylation states of KaiC, which cycle with time. The information encoded in the phosphorylation state of KaiC is then relayed to clock-controlled genes by the master transcription factor RpaA (Markson et al., 2013; Taniguchi et al., 2010) via the input/output sensor histidine kinases CikA and SasA, the phosphatase and kinase that modulate RpaA activity (Cohen, 2020). KaiA and KaiB regulate the phosphorylation state of KaiC in a negative feedback loop configuration that drives the oscillatory gene expression. In addition to the elucidation of many mechanistic details (Cohen, 2020), other investigations have provided evidence indicating that that the circadian clock in *S. elongatus* gates the cell cycle (Dong et al., 2010; Yang et al., 2010) and regulates the competence state, natural transformation being maximal when the onset of darkness coincides with the dusk circadian peak (Taton et al., 2020).

An important cyanobacterial order, *Nostocales*, consists of multicellular organisms such as *Anabaena* sp. strain PCC 7120 (henceforth *Anabaena*), in which cells are organized in a filamentary structure, with local, nearest-neighbor cell-cell coupling via septal junctions (Herrero et al., 2016). *Anabaena* bears homologs not only of the core *kai* genes of *S. elongatus* (Schmelling et al., 2017), but also of the network of genes in which they are embedded, and those coding for RpaA, as well as CikA and SasA. Whereas not much is known about the detailed molecular mechanisms behind the circadian clock in *Anabaena*, structural studies suggest that the interactions between the respective proteins are similar (Garces et al., 2004).

First insights into the dynamical behavior of clocks in *Anabaena* were obtained from bulk and DNA microarray studies that established that its circadian clock is autonomous, and that it can run freely under constant light conditions (Kushige et al., 2013). Following entrainment by two 12-h light-dark cycles, a group of genes exhibiting oscillatory behavior were identified, and the homologs of *kaiA, kaiB*, and *kaiC* genes showed low-amplitude or arrhythmic expression, in contrast to those of *S. elongatus* (Kushige et al., 2013). More recently, the collective behavior of circadian clocks in *Anabaena* filaments has been studied at the individual cell level (Arbel-Goren et al., 2021). Circadian clocks along filaments were interrogated under nitrogen-replete conditions in which all cells in the filaments were vegetative, carrying out both oxygenic photosynthesis and assimilation of a source of combined nitrogen. Under these conditions, filaments grow by binary fission of each and every cell along their length. This study found significant synchronization and spatial coherence of clock phases on the scale of filaments, evidence supporting the notion of clock coupling via cell-cell communication, and gating of the cell division by the circadian clock. Furthermore, the study confirmed the low-amplitude circadian oscillatory transcription of *kai* genes comprising the post-transcriptional core oscillatory circuit suggested by results of a bulk study (Kushige et al., 2013), and found evidence of large-amplitude oscillations of *rpaA* transcription (Arbel-Goren et al., 2021).

Under nitrogen deficient conditions, *Anabaena* fixes atmospheric nitrogen, an activity that is incompatible with the oxygen produced by photosynthesis (Flores and Herrero, 2010). The incompatibility of photosynthesis and nitrogen fixation is solved by division of labor: filaments undergo a process of development into a one-dimensional pattern consisting of single, specialized micro-oxic cells, the heterocysts, in which atmospheric nitrogen fixation takes place, separated by about 10-15 vegetative cells that fix CO_2_ photosynthetically (Corrales-Guerrero et al., 2015; Di Patti et al., 2018; Herrero et al., 2016). The genetic cascade leading to heterocyst formation is controlled by the master regulator of differentiation HetR, and involves at least two inhibitory signals related to the PatS polypeptide and the HetN protein that can be transferred from cell to cell through septal junctions. Heterocyst differentiation and the ensuing emergence of developmental patterns in *Anabaena* entail profound metabolic and morphological changes (Flores et al., 2019; Herrero et al., 2016), including some that affect cell-cell communication (Camargo et al., 2021). Results from a DNA microarray analysis of heterocyst-enriched samples have provided evidence of circadian clock activity of *kai* genes in heterocysts (Kushige et al., 2013). However, the possibility that the rhythmic transcription was indirectly induced in heterocysts by time-dependent intercellular signals from oscillators in neighboring vegetative cells could not be excluded. Moreover, the results revealed that under nitrogen deficient conditions, 39 of the 78 previously identified clock-controlled genes preserved rhythmic expression, a subset being heterocyst-specific (Kushige et al., 2013). Of note, the number of genes reported to oscillate with a circadian period in *S. elongatus* is 856, significantly larger than 78 (Markson et al., 2013).

Here we set out to study the interplay between the circadian clock and the genetic network controlling heterocyst differentiation under nitrogen deficient conditions in *Anabaena*. We tracked circadian rhythms in individual vegetative cells and heterocysts in combined nitrogen-deprived *Anabaena* filaments in real time, by following the expression from the promoter of *pecB*, a clock-controlled gene that exhibits high-amplitude oscillations (Kushige et al., 2013). This gene is part of the *pecBACEF* operon coding for the beta subunits of phycoerythrocyanin, a structural component of the phycobilisome rod that plays a major role in light harvesting for photosynthesis (Swanson et al., 1992). Our study provides evidence that nitrogen deprivation has a profound influence on the synchronization and spatial coherence of clocks along a filament, and that in addition to gating the cell cycle, the circadian clock also gates cellular differentiation.

## Results

### Discoordinated expression of a clock-controlled gene along filaments under constant light conditions

The *pecB* gene, encoding the beta subunit of phycoerythrocyanin (Swanson et al., 1992), is known to display circadian oscillations both under nitrogen-replete and nitrogen deficient conditions (Kushige et al., 2013). Expression from a chromosomal fusion of *gfp* to the 5’ region of *pecB*, denoted as *P_pecB_-gfp* (13), was followed along wild-type *Anabaena* filaments under constant light, after submitting filaments to nitrogen deprivation in BG11_0_ medium. A series of phase contrast, fluorescence and autofluorescence of photosynthetic pigments (AF) snapshots taken at maxima and minima of a number of circadian cycles is shown in Fig. 1. In contrast to the images taken right after nitrogen deprivation (t=0), in which expression along a filament was largely uniform except for small amplitude variations, at later times filaments displayed considerable heterogeneity. This heterogeneity may be due to demographic noise, or alternatively, may reflect different metabolic states in different cell stretches of the filament (Nieves-Morión et al., 2021). Expression from *P_pecB_-gfp* was visibly higher in some vegetative intervals than in others, alternating in time, and the heterogeneity in expression was spatially locked with the instantaneous pattern of heterocysts to a large extent (Fig. 1, Movie 1).

**Figure 1.**
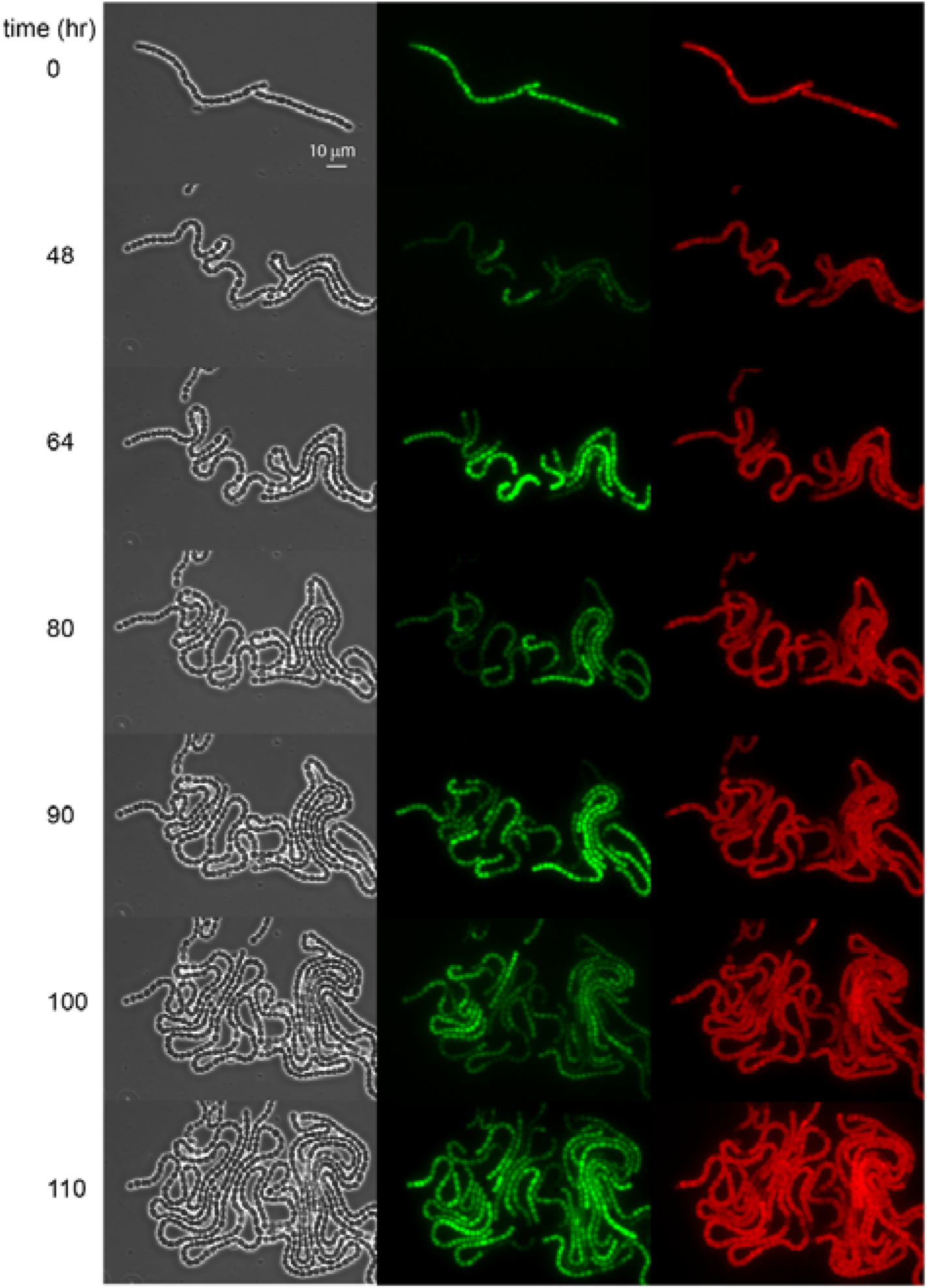
Circadian oscillations in *Anabaena* filaments under nitrogen-poor conditions. (*Left*) Phase contrast images of a filament of an *Anabaena* strain, growing under nitrogen-poor conditions. (*Middle*) GFP fluorescence in a filament of an *Anabaena* strain bearing a *P_pecB_-gfp* promoter fusion, growing under nitrogen-poor conditions. (*Right*) Autofluorescence as a function of time in *Anabaena*. Snapshots correspond to those in the left-hand micrographs and time 0 corresponds to the time at which filaments were placed under the microscope. The times at which snapshots were taken were chosen near maxima and minima of the circadian oscillations observed in GFP fluorescence intensity. For a time-lapse movie see Movie 1.

The physiological changes involved in the differentiation of a vegetative cell into a heterocyst entail alteration of cell-cell communication (Herrero et al., 2016), and altered cell-cell communication (Arévalo et al., 2021) may lead to the presence of filament cell stretches showing different metabolic states (Nieves-Morión et al., 2021). To test the notion that altered communication may affect the synchrony along a filament, we evaluated the extent of synchronization between vegetative cells belonging to different vegetative intervals, and compared it to the synchronization of vegetative cells within the same interval. To this end, we calculated the synchronization index *R* (Materials and Methods and (Garcia-Ojalvo et al., 2004)) between the two vegetative cells on either side of a given heterocyst, and compared its value to *R* calculated for two vegetative cells separated by an intervening vegetative cell (Table 1, second and third rows).

**Table 1.**
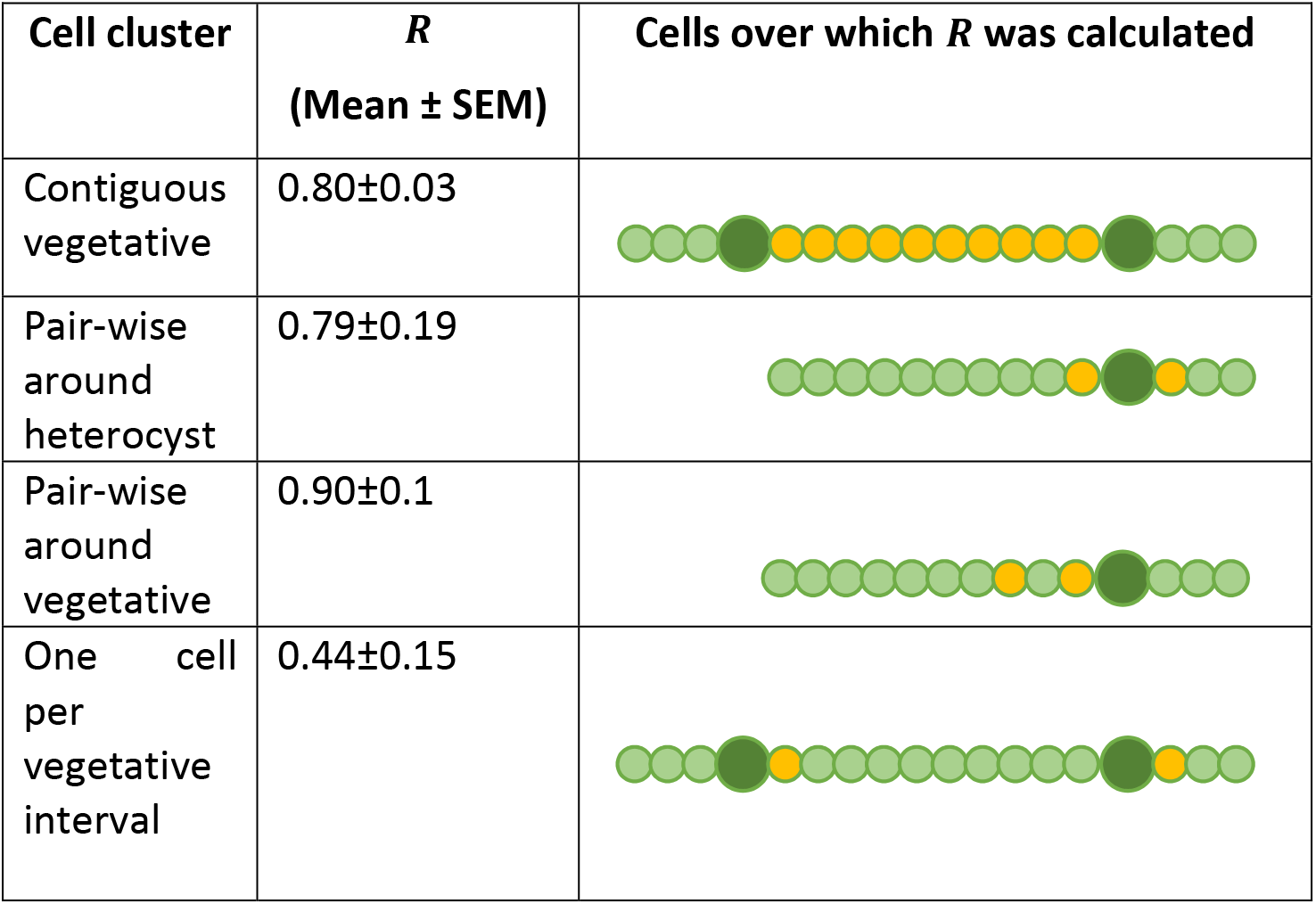
Synchronization index *R* of vegetative cells under nitrogen-poor conditions. The synchronization index *R* (Materials and methods) of vegetative cells within filaments under nitrogen-poor conditions is shown. First row: all contiguous vegetative cells within heterocysts-bounded intervals (10 cells per interval and three independent intervals). Second row: pairs of vegetative cells separated by a heterocyst (4 pairs, three independent experiments). Third row: pairs of vegetative cells separated by a single vegetative cell (4 pairs, three independent experiments). Fourth row: two vegetative cells adjacent to two heterocysts in different intervals (10 cells per filament, three independent experiments). Values of *R* represent means over the indicated number of independent experiments, and errors represent the standard errors. Yellow cells in the respective cartoons represent the vegetative cells in filaments over which *R* was calculated in each case. Dark green cells represent heterocysts and light green represent vegetative cells.

| Cell cluster | $R$<br>(Mean ± SEM) | Cells over which $R$ was calculated |
| --- | --- | --- |
| Contiguous vegetative | $0.80 \pm 0.03$ | |
| Pair-wise around heterocyst | $0.79 \pm 0.19$ | |
| Pair-wise around vegetative | $0.90 \pm 0.1$ | |
| One cell per vegetative interval | $0.44 \pm 0.15$ | |

While the mean values of *R* in both cases were comparable, the distributions of *R* values were highly skewed (Fig. 2). A comparison using a Wilcoxon-Mann-Whitney test indicates that these distributions differ significantly at the p<0.009 level. In addition, we compared the relative synchronization of vegetative intervals at the level of the whole filament. We obtained *R* =0.44±0.15 for cells from different vegetative intervals and *R* =0.80±0.03 from those within the same interval, under nitrogen-deprived conditions (Table 1). This is compared to *R* =0.59±0.06 and *R* =0.85±0.01 under nitrogen-rich conditions for separated cells in different filaments and the same filament respectively (Arbel-Goren et al., 2021). These findings indicate that, in diazotrophic filaments, oscillations in vegetative intervals are desynchronized from one another, much like cells from entirely separate filaments, but maintain a normal degree of synchrony within intervals.

**Figure 2.**
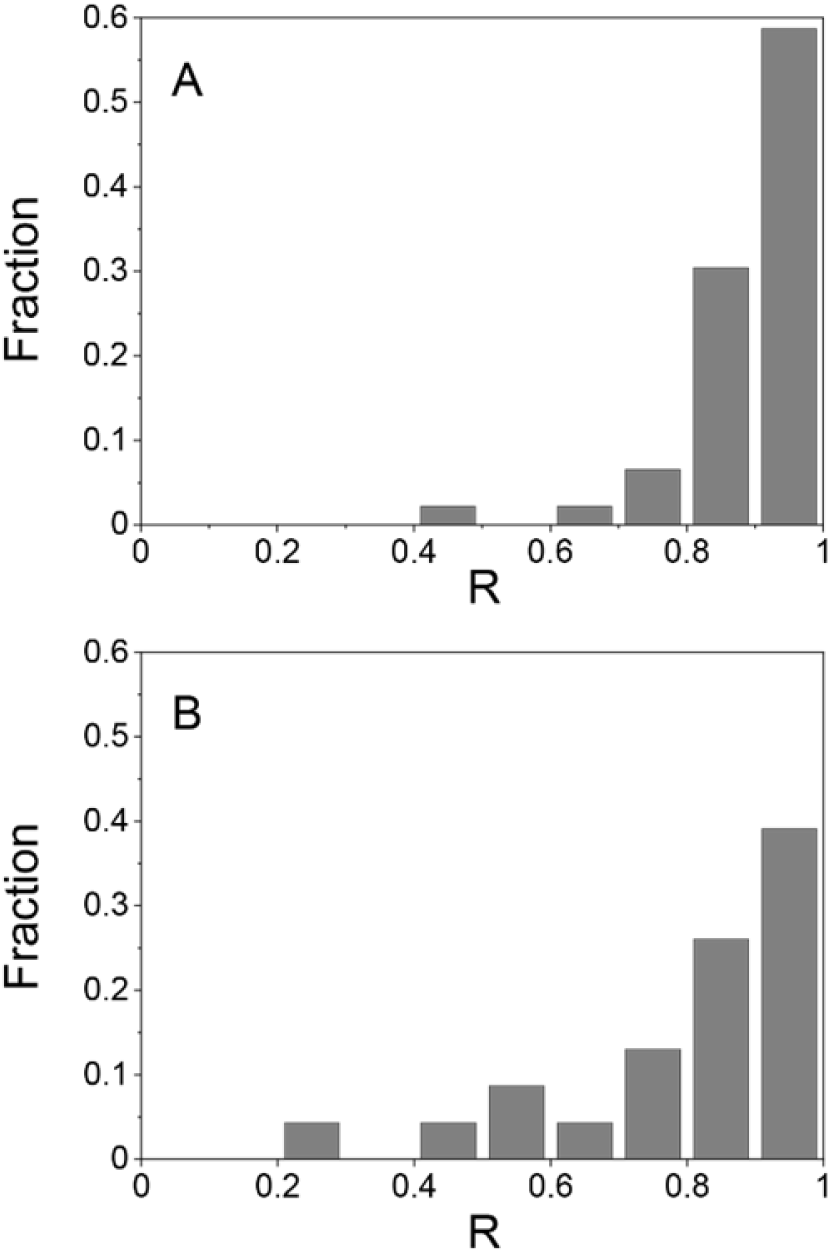
Distribution of R values between separated cells. Pair-wise values of R between vegetative cells separated by a single vegetative cell (A) or heterocyst (B). The mean and SD of the distributions from at least two independent experiments are: *R* =0.90±0.1 and *R* = 0.79 ± 0.19 respectively. A Wilcoxon-Mann-Whitney U-test indicates that these data come from continuous distributions with different medians with p=0.0086 at the 5% significance level.

### Sequential turnoff of expression in vegetative cells between consecutive heterocysts

A salient feature of oscillations in the fluorescence intensity from *P_pecB_-gfp* was the cell-cell variation in the times at which expression was turned off in vegetative cell intervals, preempting the decrease of the fluorescence intensity and the completion of a cycle. This decrease, mediated presumably both by dilution by cell growth and degradation of the GFP, resulted in a particularly wide spread of fluorescence intensity values between cells. To check whether there is coordination in expression turnoff times along a filament, we represented the fluorescence intensity in individual contiguous cells in a heterocyst-bound interval during one oscillation, color-coded according to their spatial position along the vegetative interval (Fig. 3). Notably, the fluorescence intensity during upregulation was synchronized among cells, but the decay was delayed as a function of the cell’s proximity to a heterocyst. Thus the decay in cells near the middle of a vegetative interval was most delayed. We did not detect sequential turnoff under nitrogen-replete conditions. Note also that cells near the middle of the vegetative interval appear to display higher expression.

**Figure 3.**
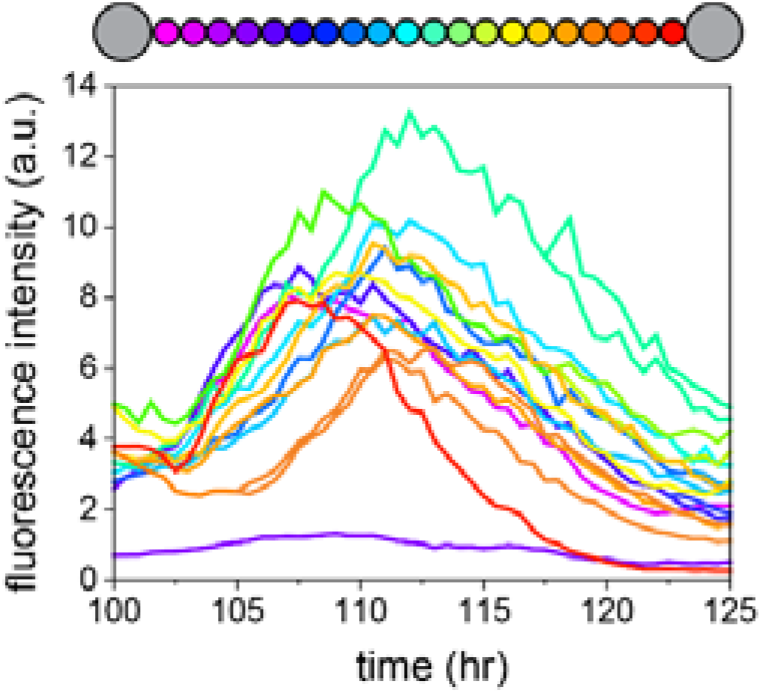
Gradient and sequential activation of fluorescence intensity from *P_pecB_-gfp* of vegetative cells within a heterocyst-bound interval. Fluorescence intensity of individual cells as a function of time over one circadian cycle. Traces are color-coded according to their position along a heterocyst-bound vegetative interval. The violet line at the bottom of the plot corresponds to the trace of one of the bounding heterocysts.

### The circadian clock gates heterocyst differentiation

To test whether the heterocyst differentiation process and circadian clocks are temporally coordinated in *Anabaena* cells, we determined the onset of the reduction of the autofluorescence of photosynthetic pigments (AF) in a cell that eventually will become a heterocyst as a temporal reference point (Foulds and Carr, 1977; Maldener et al., 1991; Wood and Haselkorn, 1980), and its phase along the cell’s circadian cycle, taking 0 and 2π to correspond to consecutive minima in the cyclic expression from *P_pecB_-gfp* (Fig. 4A). A histogram of the phases of AF intensity reduction events obtained from traces similar to those in Fig. 4A is shown in Fig. 4B. Clearly, differentiation takes place within a narrow temporal window of the circadian cycle. We conclude that the circadian clock gates heterocyst differentiation. Lastly, we note that the phase of oscillation in the heterocyst was inherited from that of the original vegetative cell.

**Figure 4.**
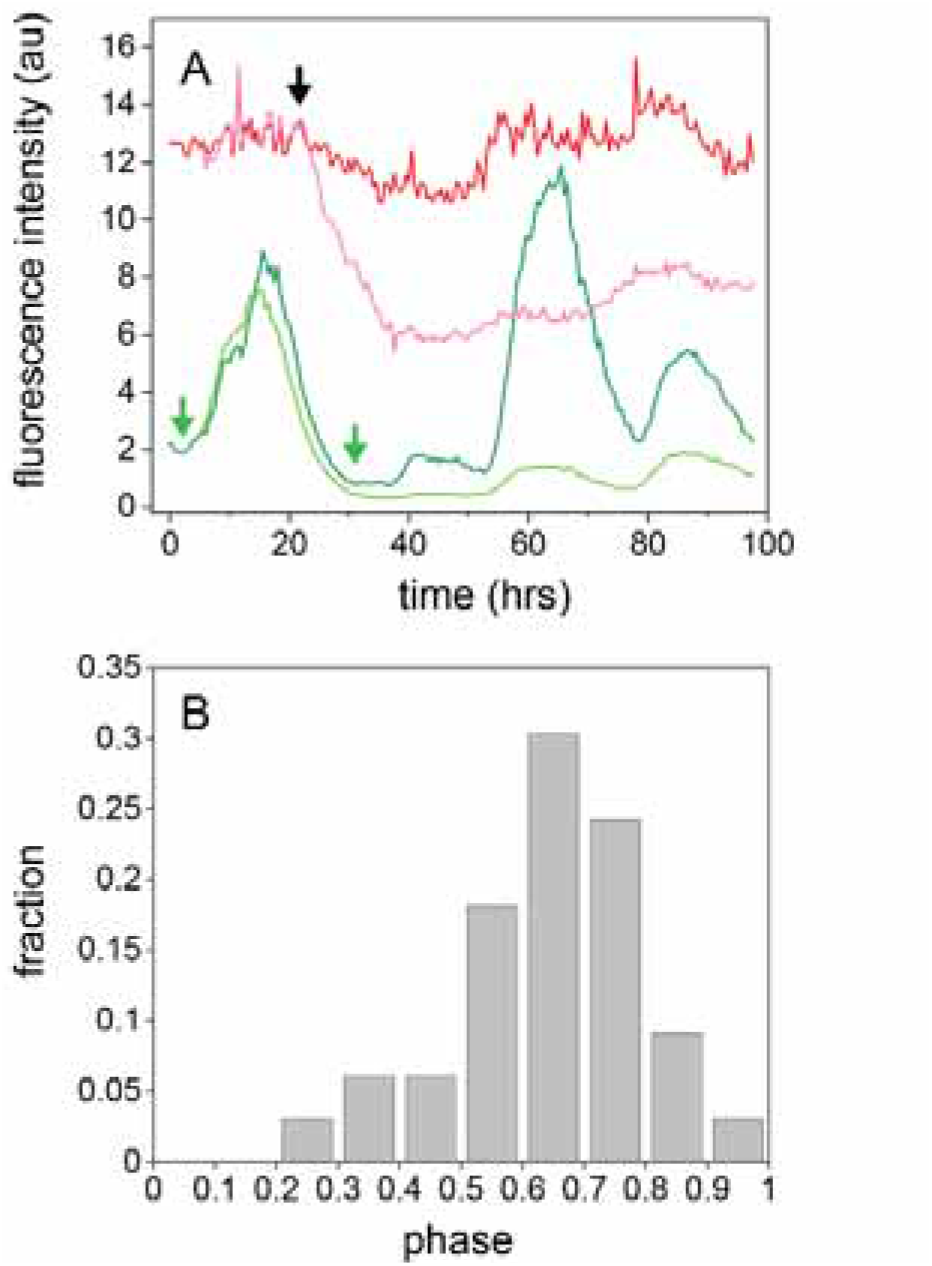
Gating of heterocyst differentiation by the circadian clock. (A) Fluorescence intensities of two sister cells bearing a *P_pecB_-gfp* fusion (green) one of which eventually becomes a heterocyst, and their respective autofluorescence intensity traces (red) as a function of time, under conditions of constant illumination. The onset in the decay of autofluorescence (AF) in the cell that becomes a heterocyst is indicated with a black arrow, whereas the positions of the circadian cycle minima on either side are indicated with green arrows. (B) Normalized histogram of the phase of onset times of autofluorescence reduction in cells that become heterocysts, with 0 and 1 denoting two consecutive minima in circadian cycles in units of 2π. Data from at least three independent experiments.

### Low amplitude circadian oscillations in heterocysts

Evidence for circadian clock activity in heterocysts was obtained previously by interrogating heterocyst-enriched bulk samples (Kushige et al., 2013). However, these experiments could not exclude the possibility that oscillations were induced in heterocysts from the transfer of time-dependent signals from neighboring vegetative cells. To test whether oscillating transcription indeed takes place in heterocysts, we followed expression from *P_pecB_-gfp* in individual heterocysts that formed after filaments were subjected to nitrogen deprivation (BG11_0_ medium). The fluorescence intensity of individual heterocysts in a typical experiment is shown in Fig. 5A. Since heterocysts formed at different times during the experiment, traces have been temporally aligned by using the onset of the decay in the autofluorescence as a temporal reference point (Foulds and Carr, 1977; Maldener et al., 1991; Wood and Haselkorn, 1980) (e.g., Fig. 5A). A comparison of these traces with those of three vegetative cells and their respective lineages (Fig. 5B) shows that the period of the oscillations in heterocysts (21.3±0.6 mean±SE n=30) is undistinguishable from that observed in vegetative cells (20.8±0.4, mean±SE n=35), and that their amplitude is about a factor of five smaller.

**Figure 5.**
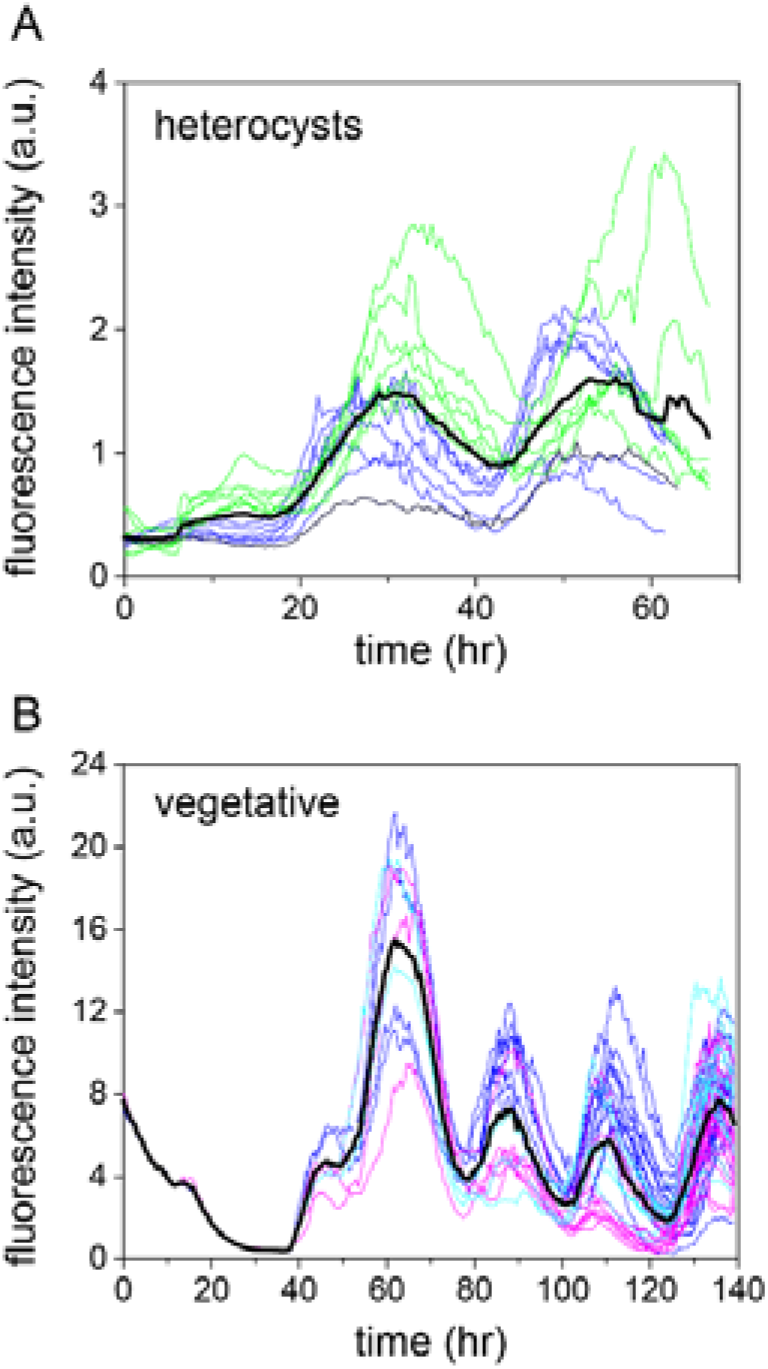
Fluorescence intensity from *P_pecB_-gfp* as a function of time in heterocysts and vegetative cell lineages. (A) Fluorescence intensity in heterocysts. Different colors correspond to data from different filaments. Traces were displaced so that the onsets of decay of autofluorescence in the vegetative cells that differentiate into heterocysts coincide. (B) Lineages of three cells each in one of three contiguous vegetative cell intervals (color-coded according to their respective interval). Thick black lines in both panels correspond to the average of all traces.

### Δ*kai* mutant filaments fail to grow fixing N_2_

To understand further the role played by the circadian clock in filament behavior under nitrogen deprivation, we studied the phenotype of filaments in which *kaiABC* genes were deleted (henceforth Δ*kaiABC* strains, Fig. S1). The growth of a Δ*kaiABC* strain was studied using four independent clones: two in which the C.K1 gene cassette was inserted in direct orientation (clones A) and two in which it was inserted in reverse orientation (clones B) with regard to the orientation of the operon. None of the clones could grow under photoautotrophic conditions in solid BG11_0_ medium, which lacks combined nitrogen (Fig. 6A). In liquid BG11_0_ medium, after an initial increase in cell mass, the four clones also failed to grow (Fig. 6B). Ten days after nitrogen deprivation, cultures from two clones bearing the inserted cassette in each of the two possible orientations were visualized by light microscopy, showing the presence of abundant cell debris and few filaments as compared to the WT (Fig. 6C). Nonetheless, heterocysts were observed in the two mutant cultures as in the WT culture, and the frequency of heterocysts was similar in the three cultures, albeit slightly higher in the mutants than in the WT at 24 h and slightly lower at 48 h (Fig. S2). Furthermore, filaments of Δ*kaiABC* strains exhibited a significantly lower production of photosynthetic pigments under nitrogen deprivation relative to the wild-type strain (Fig. 6A), precluding the detection of the decay of autofluorescence in incipient heterocysts. These observations collectively show that deletion of the *kai* genes leads to failure in diazotrophic growth while allowing heterocyst differentiation.

**Figure 6.**
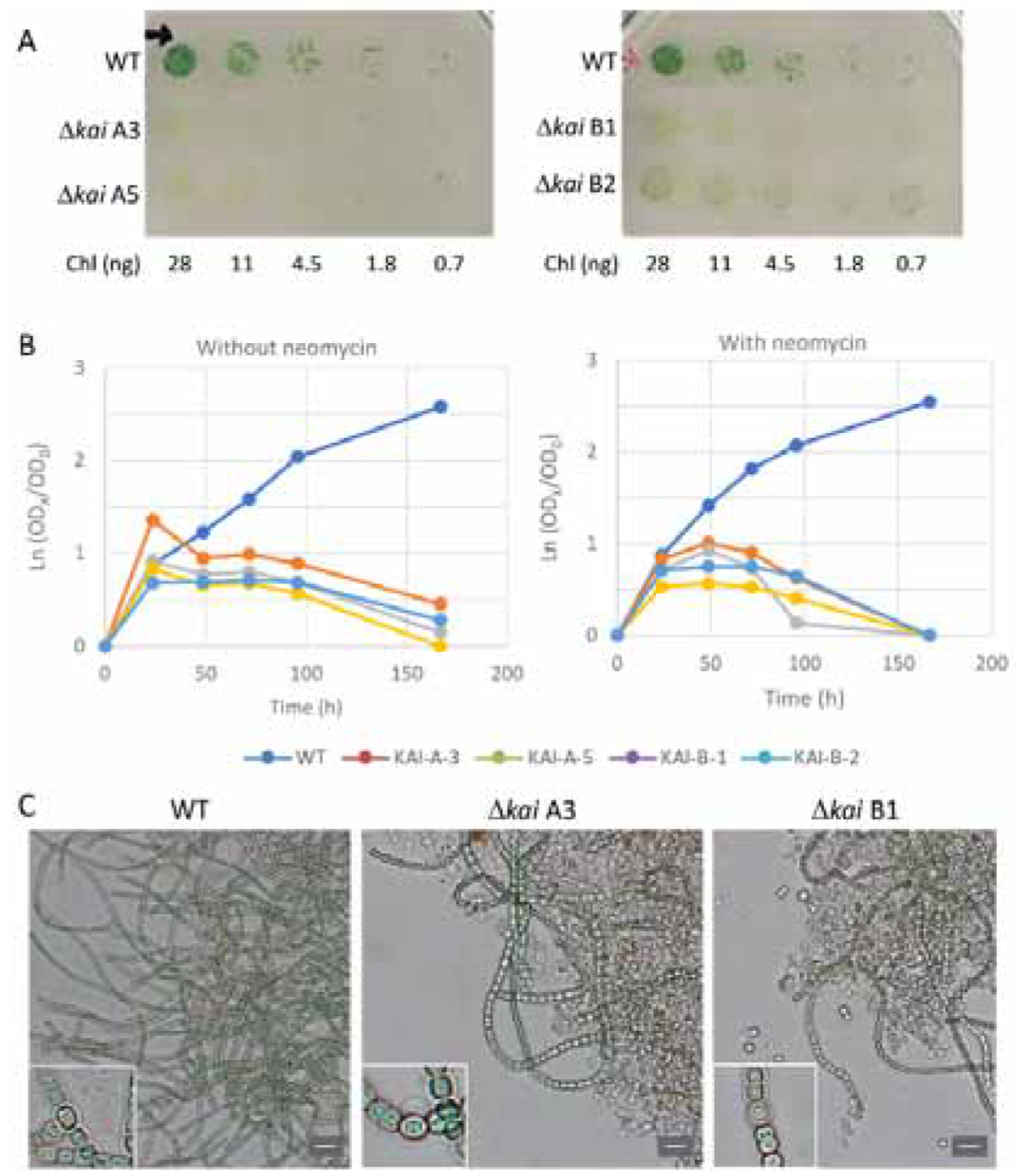
Phenotype of the Δ*kai* mutants of *Anabaena* sp. strain PCC 7210. (A) Growth tests on plates with BG11_0_ medium. The filaments were grown in BG11 medium with neomycin at 20 μg mL^-1^, washed with BG11_0_ medium (without neomycin), and incubated in BG11_0_ medium (without neomycin) for 10 days under photoautotrophic culture conditions. (B) Growth tests in liquid BG11_0_ medium without neomycin or with neomycin at 20 μg mL^-1^ for the mutants, as indicated. y axis, Ln (OD_750_ nm at time X/OD_750 nm_ at time 0); x axis, time of incubation under photoautotrophic culture conditions. For panels A and B, the wild type (WT) and two mutant clones from each orientation of the gene cassette were analyzed. (C) Bright field micrographs of the WT and Δ*kai* mutant (clones A3 and B1) after 10 days of incubation in BG11_0_ medium. Whereas the WT formed long filaments, much cell debris and only a few filaments were observed for the mutants. Size bar, 10 μm. Insets, further magnification (5x) showing the presence of heterocysts in the three strains.

### Candidate genes linking the circadian clock to the behavior of *Anabaena* under nitrogen deprivation

To shed light on the relationship between gating of differentiation and the failure in diazotrophy of Δ*kaiABC* strains on one hand, and the circadian clock on the other, we searched bioinformatically the *Anabaena* genome for a conserved signature of the RpaA-binding motif previously reported for 101 sequences in *S. elongatus* (Markson et al., 2013), taking advantage of the 96% similarity between the RpaA amino acid sequence of *S. elongatus* and *Anabaena* RpaA protein (All0129). Our search was restricted to detect the motif only within regions starting upstream to putative transcription start sites (TSSs) of genes, based on reported annotations of TSS (Mitschke et al., 2011) (motifs within a window of −500bp to +50bp relative to the TSS).

A number of genes encoding regulatory proteins were detected among 81 genes bearing putative RpaA binding sites with a FIMO q-value < 0.05 (Table S1). These include the ferric uptake regulator-related protein FurC (Alr0957), which is a protein with pleiotropic effects that affects nitrogen fixation in *Anabaena* (Sarasa-Buisan et al., 2022); the cAMP-binding transcriptional regulator Alr2325 (Suzuki et al., 2004); and the RNA polymerase sigma factor SigE, which is involved in expression of late heterocyst-specific genes (Mella-Herrera et al., 2011). Additionally, four other transcriptional regulators including a transcriptional regulator of unknown function (Alr3646) and three two-component regulators (All3822, Alr5272, Alr5069) were detected. It will be of interest to investigate in the future whether these regulators are indeed involved in circadian clock-related activities.

## Discussion

Cells in *Anabaena* filaments exhibit robust circadian rhythms under both nitrogen replete and deficient conditions. However, the single-cell observations reported here demonstrate that the behavior under both conditions differs considerably. Rather than displaying the high synchrony and spatial coherence characteristic of filaments under nitrogen-replete conditions (Arbel-Goren et al., 2021), filaments under nitrogen deprivation display noticeable differences in the phase of expression of a clock-controlled gene between different vegetative cell intervals when compared to the phase synchrony within the interval. The physiological changes involved in the differentiation of a cell into a heterocyst entail alteration of cell-cell communication breaking the symmetry of intercellular transfer: heterocysts become a sink of carbohydrates supplied by their vegetative neighbors, whereas heterocysts supply fixed nitrogen products to the neighboring vegetative cells (Herrero et al., 2016). The discoordination between vegetative filaments together with the reduced communication between vegetative cell intervals suggest that a vegetative cell interval and its delimiting heterocysts is the organismic unit in *Anabaena* under nitrogen fixing conditions, instead of the full filament as in nitrogen-replete conditions. This is further supported by the observation that heterocyst differentiation is not synchronized at the level of the whole filament under steady diazotrophic conditions.

A number of cellular processes have been reported to be regulated by circadian clocks in cyanobacteria. For example, the cell cycle is gated both in unicellular *S. elongatus* (Dong et al., 2010; Yang et al., 2010) as well as in *Anabaena* (Arbel-Goren et al., 2021). Similarly, experimental evidence supports the notion that the competence state in *S. elongatus* is regulated by the circadian clock (Taton et al., 2020). Here we found that the circadian clock also gates differentiation of vegetative cells into heterocysts, and that Δ*kaiABC* background filaments are impaired in diazotrophic growth. The fitness benefit of gating differentiation by the circadian clock is unclear, but we can surmise that the metabolic load on the cell may be minimized by avoiding differentiation during periods in which the cell is engaged in other processes that may compete with it. This notion is consistent with our observation that most differentiation events occur primarily when cell division events are infrequent (Arbel-Goren et al., 2021), and with the possibility that cell division of mother cells is not an essential requirement for heterocyst differentiation after nitrogen step-down (Asai et al., 2009). This possibility is under current discussion. The fact that the phase of the clock in a heterocyst is inherited from the progenitor cell, when compared to the phase of the clock of the progenitor’s sister cell, indicates that while the clock gates differentiation, the clock itself is rather insensitive to the differentiation process, despite the attendant metabolic changes involved in the differentiation process.

Interestingly, the absence of a clock does not prevent differentiation of (non-functional) heterocysts. A possible clue that may point to a mechanism behind gating of differentiation by the circadian clock is furnished by the 5-10-fold reduction in the levels of *pecB* transcription in heterocysts relative to vegetative cells. Oscillations are transmitted from the core clock to the *pecBACEF* operon most probably by the master transcription factor RpaA (Arbel-Goren et al., 2021). Here we found that upstream of *pecB* there are two putative RpaA binding sites (p value=1.5×10^-5^, Table S1). Since neither the abundance of RpaA nor its transcription decrease as a result of nitrogen deprivation (Camargo et al., 2021; Zhang et al., 2021), lower levels of *pecBACEF* transcription may be effected by a reduction in the levels of the active phosphorylated form (RpaA~P), which may be mediated by SasA and CikA in *Anabaena* as in *S. elongatus* (Gutu and O’Shea, 2013). SasA is regulated in *S. elongatus* by the phycobilisome-associated B protein (RpaB), which is involved in the integration of temporal and environmental information and stress (Espinosa et al., 2015). The conservation of the corresponding genes lends support to these notions (Schmelling et al., 2017). Together, these considerations suggest that a link between the circadian clock and the heterocyst differentiation network may be gleaned from the set of genes whose expression is regulated by RpaA~P.

Our observation of circadian oscillations in the transcriptional activity of *pecB* in individual heterocysts, together with the inheritance of the phase of oscillation from the primordial vegetative cell lead us to posit that the circadian clock continues to function in the heterocyst, and that rhythmic transcription in the heterocyst is not induced indirectly by time-dependent intercellular signals from clocks in neighboring vegetative cells. While photosystem II (PSII) is altered in heterocysts (Magnuson and Cardona, 2016), PSI continues to function (Magnuson and Cardona, 2016), and the oscillatory behavior of transcriptional activity of *pecB*, even if smaller, suggests that the circadian clock may modulate photosynthetic activity.

The sequential turnoff of gene expression according to position along a vegetative cell interval is characterized by timescales that are considerably longer that those typical of intercellular transport of metabolites (Nürnberg et al., 2015), which help maintain filaments in homeostasis. The sequential turnoff here observed is reminiscent of the waves of gene expression measured across different parts of *Arabidopsis thaliana* plants (Endo, 2016; Gould et al., 2018). Nonetheless, the signals coupling clocks are unknown as they are in higher plants (Greenwood and Locke, 2020), and remain to be elucidated.

The gating of differentiation by the circadian clock and failure of diazotrophy of Δ*kaiABC* strains led us to investigate the relationship between these two processes using bioinformatics methodologies. The high conservation of circadian clock components among cyanobacteria (Schmelling et al., 2017), and in particular the high similarity between the protein sequences of RpaA in *Anabaena* and in *S. elongatus*, suggested that RpaA function is conserved as a master clock output regulator. Therefore, we looked for the presence of RpaA putative binding sites upstream of *Anabaena* genes, with low FIMO q-values (i.e., high significance). We found that some of the ChIP-validated genes in *S. elongatus* (Markson et al., 2013), have orthologs in *Anabaena* and have putative binding sites of RpaA. Together, these findings support the notion that RpaA may play a functional role as master regulator of clock outputs in *Anabaena* as in *S. elongatus*. However, we have also identified in *Anabaena* putative RpaA controlled genes with specific roles in heterocyst function, which would explain the lack of heterocyst activity in Δ*kaiABC* strains. In summary, our work revealed that in *Anabaena* the circadian clock is further necessary to confront nitrogen stress.

## Materials and Methods

### Strains

Strains bearing a chromosomally encoded *P_pecB_-gfp* were obtained by conjugation with the *Anabaena* sp. (also known as *Nostoc* sp.) PCC 7120 wild-type background and with a Δ*kaiABC* background, in which the *kaiABC* genes were deleted (Arbel-Goren et al., 2021), as recipients.

### Culture conditions

Strains and derived strains were grown photoautotrophically in BG11 medium containing NaNO_3_, supplemented with 20 mM HEPES (pH 7.5) with shaking at 180 rpm, at 30 °C, as described previously (Corrales-Guerrero et al., 2014, 2013). Growth took place under constant illumination (10 μmol m^-2^s^-1^) of photons (spectrum centered at 450 nm) from a cool-white LED array. When required, streptomycin sulfate (Sm), and spectinomycin dihydrochloride pentahydrate (Sp) were added to the media at final concentrations of 2 μg/mL for liquid and 5 μg/mL for solid media (1% Difco agar); neomycin sulfate (Nm) was added at 10 and 25 μg/mL, respectively. The densities of the cultures were adjusted so as to have a chlorophyll *a* content of 2-4 μg/mL 24 h prior to the experiment, following published procedures (Di Patti et al., 2018). For time lapse measurements, filaments in cultures were harvested and concentrated 50 fold.

### Samples for time-lapse microscopy

Strains were grown as described previously (Di Patti et al., 2018). When required, antibiotics, streptomycin sulfate (Sm) and spectinomycin dihydrochloride pentahydrate (Sp), were added to the media, at final concentrations of 2 μg/mL for liquid and 5 μg/mL for solid media. The densities of the cultures, grown under an external LED array (15 μmol m^-2^s^-1^) for about five days, were adjusted so as to have a chlorophyll *a* content of 2-4 μg/mL, 24 h prior to the experiment following published procedures (Di Patti et al., 2018). For time-lapse, single-cell measurements of *Anabaena*, 5 μL of culture concentrated 100-fold were pipetted onto an agarose low-melting gel pad (1.5%) in BG11 medium containing NaNO_3_ and 10 mM NaHCO_3_, which was placed on a microscope slide. The pad with the cells was then covered with a #0 mm coverslip and then placed on the microscope at 30 °C. The cells grew under light from both an external LED array (15 μmol m^-2^s^-1^) and tungsten halogen light (10 μmol m^-2^s^-1^), 3000K colour). Under these illumination conditions, the doubling time of cells is similar to that in bulk cultures (Di Patti et al., 2018). The change in illumination conditions when transferring cells from bulk cultures to the microscope results in high synchronization within filaments. Images of about ten different fields of view were taken every 30 min on a Nikon Eclipse Ti-E microscope controlled by the NIS-Elements software using a 60 N.A 1.40 oil immersion phase contrast objective lens (Nikon plan-apochromat 60 1.40) and an Andor iXon X3 EMCCD camera. Focus was maintained throughout the experiment using a Perfect Focus System (Nikon). All the filters used are from Chroma. The filters used were ET480/40X for excitation, T510 as dichroic mirror, ET535/50M for emission (GFP set), ET500/20x for excitation, T515lp as dichroic mirror, and ET535/30m for emission (EYFP set), and ET430/24x for excitation, 505dcxt as dichroic mirror, and HQ600lp for emission (chlorophyll set). Samples were excited with a pE-2 fluorescence LED illumination system (CoolLED).

### Image segmentation

All image processing and data analysis was carried out using Matlab (MathWorks). Filament and individual cell recognition was performed on phase contrast images using an algorithm developed in our laboratory. The program’s segmentation was checked in all experiments and corrected manually for errors in recognition. The total fluorescence from GFP and chlorophyll *a* (autofluorescence) channels of each cell, as well as the cell area, were obtained as output for further statistical analysis.

### Analysis of synchronization along filaments

Synchronization was measured by the order parameter (Garcia-Ojalvo et al., 2004):

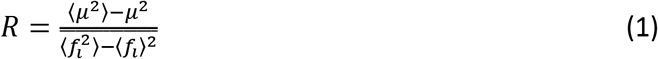

where 〈·〉 denotes a time average, 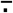 indicates an average over all cells, and *μ* denotes the average of the fluorescence intensity of each cell *f_i_*. Hence *R* is defined as the ratio of the standard deviation of *μ*(*t*) to the standard deviation of *f_i_*, averaged over all cells.

For measurement of synchronization within the same filament, groups of 8-11 cells were chosen, whether separated or contiguous (sharing a common ancestor as determined from a lineage analysis). For evaluation of inter-filament synchronization, one cell per filament was chosen randomly in different fields of view. R was then calculated and this procedure was repeated for different choices of cells, at least three times for each experiment. All the evaluations of R were carried out over a full period of oscillation, in either one of the first two oscillations, except for the Δ*kaiABC* background, for which R was calculated for an interval of 24 hours, during which other strains display the first full oscillation. The final result comprises the mean of at least three independent repeats, in at least two independent experiments. Errors in the quoted values of R therefore represent standard errors (SEM). Statistical analyses were performed in Matlab using Mann-Whitney’s U-test.

### Identification of RpaA binding motifs in the *Anabaena* genome

Motif scanning was done using FIMO tool from the MEME Suite (v5.4.1) (Grant et al., 2011), using a previously reported DNA Position-specific probability matrix of the RpaA binding motif in *S. elongatus* (Markson et al., 2013), scanning both strands and reporting a minimal match p-value of 10^-4^. The results were restricted to sequences within a window of - 500bp to +50bp relative to the TSS, based on previously reported *Anabaena* TSS annotations (Mitschke et al., 2011). Assignment of resulting motifs to genes was based on annotations of valid gene names (Mitschke et al., 2011), as well as reported annotations of early and late differentiation genes (cluster #6 and Cluster #4 genes (Brenes-Álvarez et al., 2019)) and (Kushige et al., 2013). Protein sequences coded by *Anabaena* genes, which harbor a putative RpaA motif, were compared with genes reported to bind RpaA in a ChIP experiment in *S. elongatus* (Markson et al., 2013). Thus, protein sequences of 89 reported PCC 7942 genes were compared using BLASTP to *Anabaena* proteins, reporting hits with E-value =<0.005 and >36% sequence similarity (Table S2).

## Acknowledgements

Work in Rehovot was supported by the Minerva Stiftung http://www.minerva.mpg.de/ to JS. JS is the incumbent of a Siegried and Irma Ullman Professorial Chair. Work in Seville was supported by grant no. PID2020-118595GB-100 from Agencia Estatal de Investigación, Spain, and the European Regional Development Fund to AH and EF.

## Data availability

Source data files and Matlab code have been deposited in Dryad (DOI https://datadryad.org/stash/share/HOU6G8tz9ugDga2XO_GmaPfNTgfCAYMu8DbI2jFqxUQ).

## Supporting files

**Movie 1.** Circadian oscillations in *Anabaena* filaments under nitrogen-poor conditions as a function of time.

**Figure S1.**
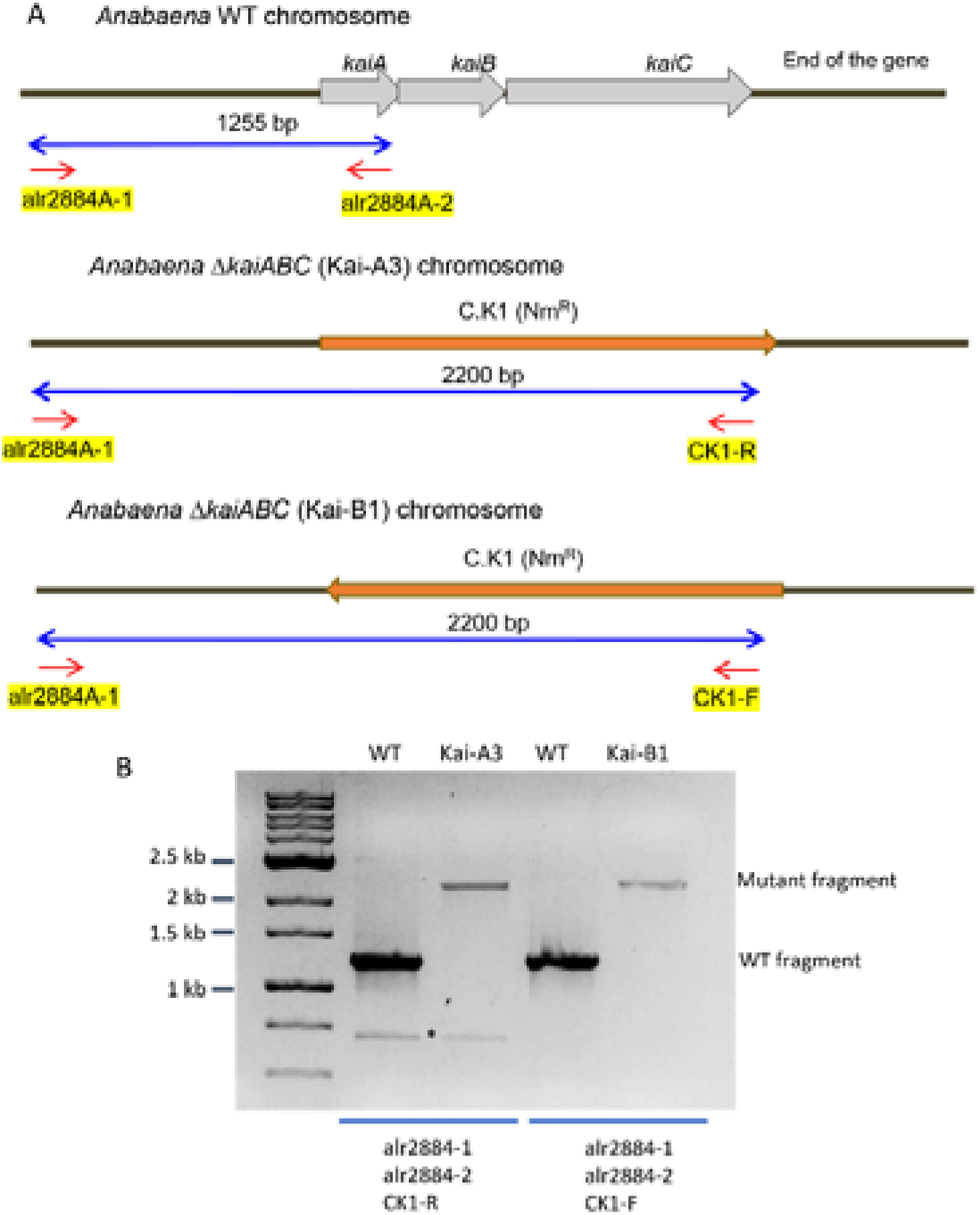
Genetic structure in the Δ*kaiABC* mutants. The Δ*kaiABC* mutants were re-isolated following the procedure described in Arbel-Goren *et al.*, 2021, but now the gene-cassette, C.K1, was inserted in both orientations (direct orientation, mutants A; opposite orientation, mutants B). (A) The genetic structure in the *kai* genomic region is shown for the wild type (top scheme), a mutant with the gene cassette in direct orientation (middle) and a mutant with the gene cassette in opposite orientation (bottom). Oligonucleotide primers used in PCR analysis are indicated. (B) PCR analysis with genomic DNA isolated from the wild type or the A3 and B1 *kai* mutants grown in BG11 medium (with neomycin at 20 μg mL^-1^ for the mutants) and incubated for 48 h in BG11_0_ medium (without neomycin). Three primers were added to each reaction, as indicated, resulting in amplification of only WT DNA fragments in the wild type and only mutant fragments in the mutants, indicating segregation of the mutant chromosomes. *, non-specific amplification product.

**Figure S2.**
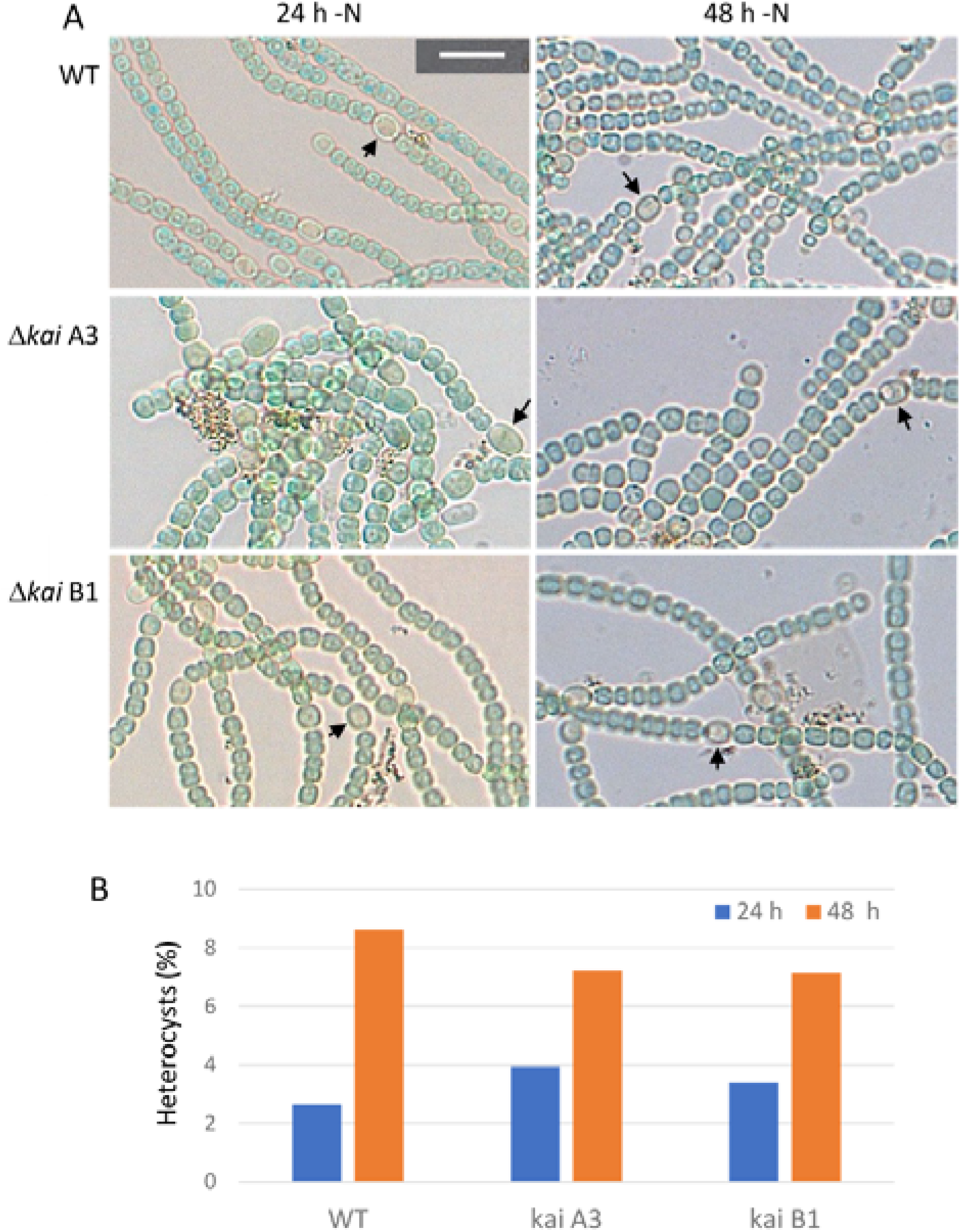
Heterocyst formation in the Δ*kai* mutants of *Anabaena* sp. strain PCC 7210. The strains were grown in BG11 medium (in the presence of neomycin at 20 μg mL^-1^ for the mutants), washed with BG11_0_ medium and incubated in liquid BG11_0_ medium (without antibiotic) under photoautotrophic culture conditions for 24 and 48 h, respectively. (A) Examples of filaments showing the presence of heterocysts (some indicated by black arrows). Size bar, 10 μm; same magnification in all the micrographs. (B) Heterocysts as percentage of total number of cells in the three strains after 24 or 48 h of incubation in BG11_0_ medium. Total number of cells counted: 1300 to 1500 in the 24-h samples; 1000 to 1100 in the 48-h samples.

**Table S1**. RpaA putative binding motifs identified in *Anabaena* sp. strain PCC 7120 including FIMO statistics and additional annotations.

**Table S2.** BLASTP analysis of *S. elongatus* orthologous proteins to *Anabaena* sp. strain PCC 7120.

